# Prostaglandin E_2_ promotes intestinal inflammation via inhibiting microbiota-dependent regulatory T cells

**DOI:** 10.1101/2020.07.12.199513

**Authors:** Siobhan Crittenden, Marie Goepp, Jolinda Pollock, Calum T. Robb, Danielle J. Smyth, You Zhou, Robert Andrews, Victoria Tyrrell, Alexander Adima, Richard A. O’Connor, Luke Davies, Xue-Feng Li, Hatti X. Yao, Gwo-Tzer Ho, Xiaozhong Zheng, Amil Mair, Sonja Vermeren, Bin-Zhi Qian, Damian J. Mole, Jürgen K.J. Schwarze, Richard M. Breyer, Mark J. Arends, Valerie B. O’Donnell, John P. Iredale, Stephen M. Anderton, Shuh Narumiya, Rick M. Maizels, Adriano G. Rossi, Sarah E. Howie, Chengcan Yao

**Affiliations:** Centre for Inflammation Research, Queen’s Medical Research Institute, The University of Edinburgh, Edinburgh EH16 4TJ, UK; SRUC Veterinary Services, Scotland’s Rural College, Easter Bush Estate, EH26 0PZ, UK; Wellcome Centre for Molecular Parasitology, Institute for Infection, Immunity and Inflammation, University of Glasgow, Glasgow G12 8TA, UK; Systems Immunity University Research Institute, and Division of Infection and Immunity, Cardiff University, Cardiff CF14 4XN, UK; MRC Centre for Reproductive Health, Queen’s Medical Research Institute, The University of Edinburgh, Edinburgh EH16 4TJ, UK; Department of Veterans Affairs, Tennessee Valley Health Authority, and Department of Medicine, Vanderbilt University Medical Center, Nashville, Tennessee, USA; Division of Pathology, Cancer Research UK Edinburgh Centre, The University of Edinburgh, Institute of Genetics & Molecular Medicine, Edinburgh EH4 2XR, UK; Senate House, University of Bristol, Bristol BS8 1TH, UK; Alliance Laboratory for Advanced Medical Research and Department of Drug Discovery Medicine, Medical Innovation Center, Kyoto University Graduate School of Medicine, Kyoto 606-8507, Japan

## Abstract

The gut microbiota fundamentally regulates intestinal homeostasis and disease partially through mechanisms that involve modulation of regulatory T cells (Tregs), yet how the microbiota-Treg crosstalk is physiologically controlled is incompletely defined. Here, we report that prostaglandin E_2_ (PGE_2_), a well-known mediator of inflammation, inhibits mucosal Tregs in a manner depending on the gut microbiota. PGE_2_ through its receptor EP4 diminishes Treg-favorable commensal microbiota. Transfer of the gut microbiota that was modified by PGE_2_-EP4 signaling modulates mucosal Treg responses and exacerbates intestinal inflammation. Mechanistically, PGE_2_-modified microbiota regulates intestinal mononuclear phagocytes and type I interferon signaling. Depletion of mononuclear phagocytes or deficiency of type I interferon receptor contracts PGE_2_-dependent Treg inhibition. Taken together, our findings provide emergent evidence that PGE_2_-mediated disruption of microbiota-Treg communication fosters intestinal inflammation.

## Introduction

Inflammatory bowel disease (IBD) is a chronic inflammatory disorder of the intestine that causes abdominal pain, diarrhea, bleeding and increased risk of intestinal cancer. There are two main subtypes of IBD, i.e., Crohn’s disease (CD) and ulcerative colitis. Multiple factors including lifestyle (e.g. smoking, diet, medications, psychological state), environmental risk factors (e.g. infections, air pollution), genetic and epigenetic alterations, and host immune functions can potentially trigger the development and progression of IBD^1,2^. The gut microbiota plays a critical role in maintaining health of the host. Dysfunction of this symbiosis may result in development of various human diseases like IBD, metabolic syndrome, infections, allergy and cancer ^3,4^. Interplay between the host and gut microbiota controls intestinal homeostasis and inflammatory responses through mechanisms that involve modulation of gut-resident regulatory T cells (Tregs) which express the transcription factor, forkhead box P3 (Foxp3) ^5–8^. Dysregulation of intestinal Tregs is implicated in the pathogenesis of IBD ^9^. Microbial antigens, metabolites (e.g., vitamins and short chain fatty acids (SCFAs)) and signaling molecules released during tissue damage (e.g., alarmins) contribute to the induction of distinct intestinal Treg subsets, that play critical roles in intestinal homeostasis and control mucosal inflammation ^10–13^. However, the mechanisms that negatively regulate microbiota-Treg crosstalk for IBD pathogenesis are incompletely studied.

Prostaglandins (PGs) are bioactive lipid mediators that are generated from arachidonic acid via cyclooxygenases (COXs) and specific PG synthases ^14^. The PG family has 5 members including prostaglandin E_2_ (PGE_2_), PGD_2_, PGF_2α_, PGI_2_ and thromboxane A_2_ (TXA_2_). PGs signal in an autocrine and/or paracrine manner through their distinct G-protein coupled receptors including PGE_2_ receptor EP1-4, PGD_2_ receptor DP1-2, PGF_2α_ receptor FP, PGI_2_ receptor IP and TXA_2_ receptor TP. PGE_2_ is present in most tissues at biologically functional nanomolar levels in the steady state, and its levels are increased at the sites of inflammation such as multiple sclerosis, arthritis, psoriasis, IBD etc ^14–16^. Non-steroidal anti-inflammatory drugs (NSAIDs), like aspirin and indomethacin, are widely used to reduce pain, fever and inflammation by inhibiting COX activities and therefore decreasing PG production. However, NSAIDs are generally avoided for individuals who have gut conditions due to the gastrointestinal adverse effects ^17^. This is because PGE_2_ plays critical roles in maintaining the gut epithelium, protecting against acute damage and facilitating regeneration after injury through actions on various cell types including macrophages, epithelial, stromal and innate lymphoid cells ^18–21^. On the other side, numerous epidemiological studies have suggested that use of NSAIDs is associated with reduced risks of colorectal cancer in the context of chronic inflammation ^22,23^.

Genome-wide association studies (GWAS) have revealed that polymorphisms in the *PTGER4* gene (encoding human PGE_2_ receptor EP4) are associated with over-expression of EP4 and a more severe disease phenotype in patients with IBD ^24–26^. Moreover, variants in the *PTGER4* gene exert a significant association with CD, in third place among all susceptible genetic loci after variants in *NOD2* and *IL23R* genes ^27^. These findings raised a possibility that PGE_2_-EP4 signaling may also participate in the pathogenesis of intestinal inflammation despite its protective actions on the epithelium. We and others have recently found that PGE_2_ plays key roles in immune-related chronic inflammatory diseases in rodents and humans, for example multiple sclerosis, rheumatoid arthritis and inflammatory skin disorders, through promoting IFN-γ-producing type 1 helper T (Th1) cells and IL-17-producing type 17 helper T (Th17) cells by induction of key cytokine receptors IL-12Rβ2 and IL-23R, respectively ^28–31^. Yet, the action of PGE_2_ on Treg responses, especially in the intestine, remains unknown. In this study, we investigated the roles for endogenous PGE_2_ in regulation of mucosal Treg responses and intestinal inflammation. We demonstrate that PGE_2_ down-regulates intestinal Treg responses by affecting the composition of the gut microbiota and modulating the function of mononuclear phagocytes (MNPs).

## Results

### Production of PGs and expression of their receptors in the intestine

Firstly, we examined whether intestine tissues physiologically produce PGs in the steady state. We administered naïve wild-type (WT) C56BL/6 mice with a pan-COX inhibitor, indomethacin, at the dose of 5 mg/kg/d (equaling to ~30 mg/d for an adult human weighing 75 kg) or vehicle (0.5% ethanol) in drinking water. Intestine tissues were collected on day 5 for measuring levels of PGs by lipidomic analysis. We observed that healthy small and large intestines in control mice produced considerable levels (i.e. hundreds to thousands nanogram per gram dry tissue) of PGs including PGE_2_, PGD_2_, PGF_2α_, TXA2 metabolite (i.e., TXB_2_) and PGI_2_ metabolite (i.e., 6-keto PGF_1α_) (**Fig. 1a and Supplementary Fig. 1**). Small intestine produced more PGD_2_ and TXB_2_, but less PGF_2α_ and 6-keto PGF_1α_, than the colon, while both intestinal tissues had comparable levels of PGE_2_ and its metabolite (13,14-dihydro-15-keto PGE_2_) or epimer (8-iso PGE_2_) (**Fig. 1a)**. Administration of indomethacin markedly inhibited production of all prostaglandins (e.g. PGE_2_, PGD_2_, PGF_2α_) and their inactive metabolites, i.e., 15-keto PGE_2_, 15-keto PGD_2_, 15-keto PGF_2α_, 6-keto-PGF_1α_ and TXB_2_ in both small and large intestines (**Fig. 1a**). For example, indomethacin reduced PGE_2_ levels by ~96% (from 2,628 to 105 ng/g dry tissue) in small intestines and by ~99% (from 2,789 to 29 ng/g dry tissue) in colons under the physiological condition (**Fig. 1a**). Re-analysis of public datasets^32^ revealed that both mouse and human intestines express EP4 genes (i.e. *Ptger4* and *PTGER4*, respectively) at the levels remarkedly higher than other PG receptors (**Fig. 1b, c**), indicating that the PGE_2_-EP4 pathway may play a more important role than other PG signaling in the intestine.

**Figure 1.**
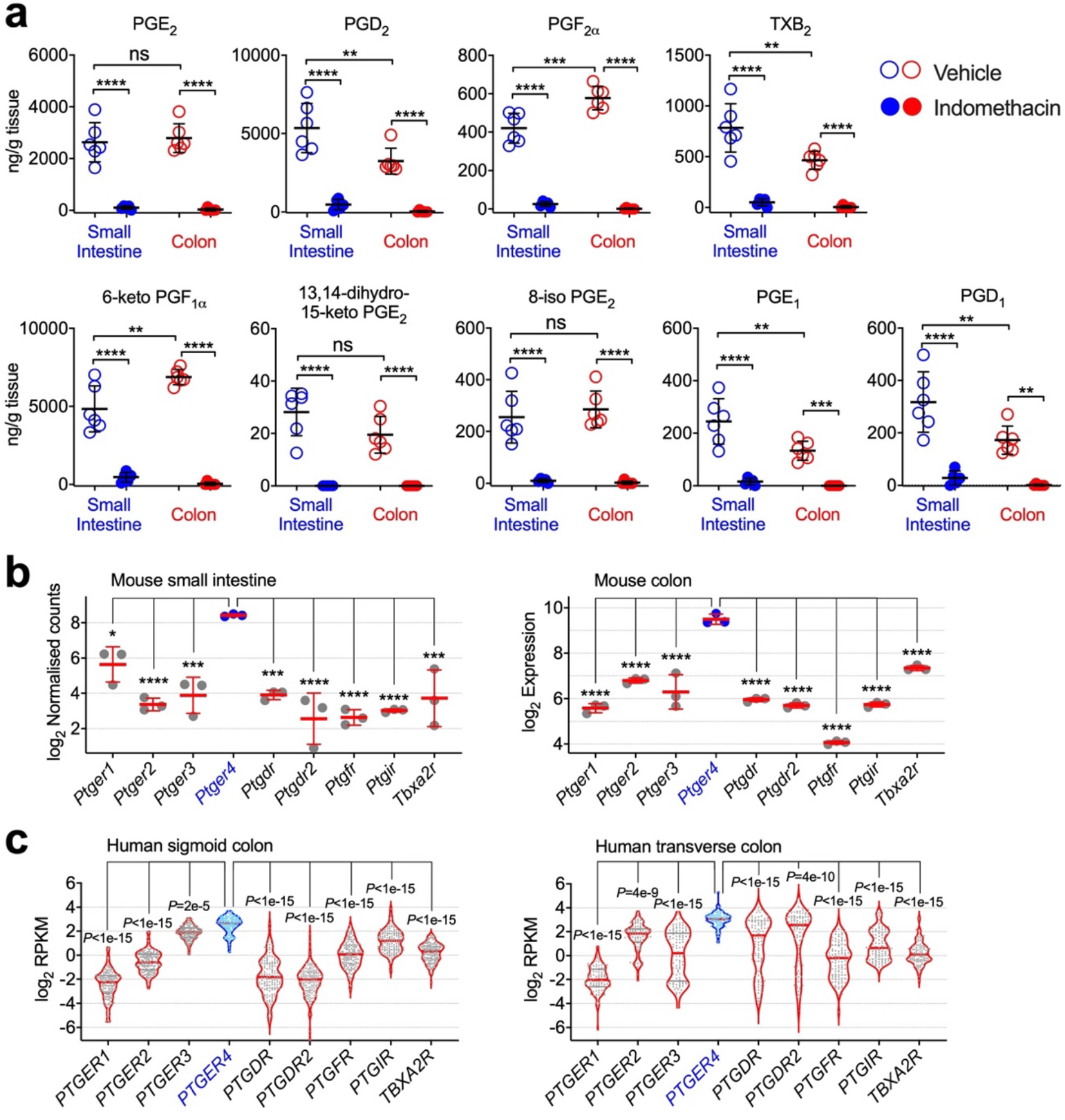
Expression of prostaglandins (PGs) and their receptors in the intestine. **(a)** Levels of PGs and their metabolites in small intestines and colons from mice administered with indomethacin or vehicle control in drinking water for 5 days (n=6 each group) as measured by lipidomic analysis. (**b**) Gene expression of PG receptors in mouse small and large intestines. Microarray gene expression data of normal C57BL/6 mouse colons (n=3) were retrieved from the Gene Expression Omnibus (GEO) dataset GSE31106. RNA-seq data of normal C57BL/6 mouse small intestines (n=3) were retrieved from the GEO dataset GSE97371. (**c)** Gene expression of PG receptors in human sigmoid (n=149) and transverse (n=104) colon biopsy samples of healthy individuals. RNA-seq data were downloaded from the Genotype-Tissue Expression (GTEx) project database and analyzed using Python 3.7.0. Each scatter dot plot in bar graphs represents data from one mouse (**a, b**) or individual (**c**). Data shown as mean ± SD (**a, b**) or presented as violin bars with scatter plots (**c**) are analysed by ANOVA with post-hoc Holm-Sidak’s multiple comparisons test. \**P*<0.05, \*\**P*<0.01, \*\*\**P*<0.001, \*\*\*\**P*<0.0001, and ns=not significant.

### Suppression of intestinal Tregs by PGE_2_-EP4 signaling in the steady state

To test whether endogenous PGs regulate intestinal Tregs in the steady state, we administered WT mice with indomethacin in drinking water and analyzed Tregs in various organs including colons, small intestines, mesenteric lymph nodes (mLN) and spleens. Indomethacin treatment increased Treg accumulation in all of these tissues, but the effect of indomethacin was greater in intestinal lamina propria (LP) (by ~2-fold in colon and small intestine) than that in mLN and spleen (both by ~1.4-fold) (**Fig. 2a, b**). Furthermore, indomethacin significantly increased mean fluorescence intensity (MFI) of Foxp3 among Foxp3^+^ Tregs in the colon (**Fig. 2b**), suggesting that inhibition of endogenous PG biosynthesis not only increases intestinal Treg cell frequencies but also enhances Foxp3 expression at the single cell level. To exclude the possibility that indomethacin-dependent increase in intestinal Tregs may be due to its destructive effects on the gut epithelium, we administered mice with indomethacin at the doses of 1-2 mg/kg body weight/day which is the equivalent of ~6-12 mg/d for adults weighing 75 kg, levels known not to induce intestinal damage in human. Low doses of indomethacin still increased intestinal Foxp3^+^ Tregs (**Supplementary Fig. 2**). As indomethacin inhibits all PG production, we next examined whether PGE_2_ and its receptor EP4 regulate intestinal Tregs. We co-administered mice with indomethacin together with a selective EP2 agonist (butaprost) or a selective EP4 agonists (L-902,688), and found that increase in colonic Tregs by indomethacin was completely prevented by the EP4 agonist in colons and mLNs, whilst the EP2 agonist only reduced Tregs in mLNs (**Fig. 2a, b**), indicating that PGE_2_ suppresses colonic Treg accumulation mainly through EP4.

**Figure 2.**
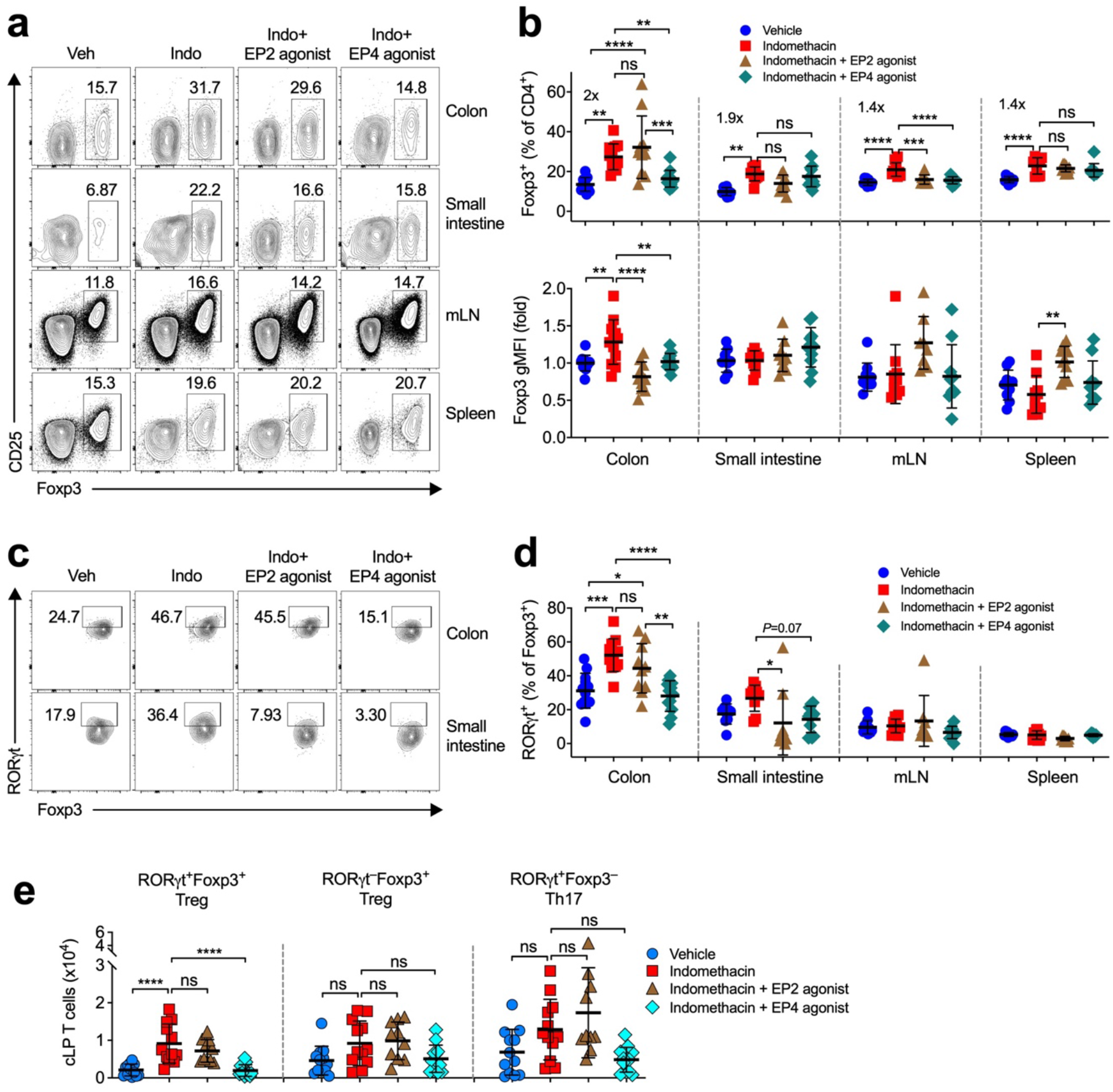
PGE_2_-EP4 signaling inhibits intestinal Tregs in the steady state. **(a)** Representative flow cytometry plots of Foxp3^+^ Tregs gated on live CD45^+^CD3^+^CD4^+^ T cells in colon and small intestinal lamina propria, mesenteric lymph node (mLN) and spleen from C57BL/6 mice that were treated with vehicle or indomethacin in drinking water and injected intraperitoneally with an EP2 agonist, Butaprost, or an EP4 agonist, L-902,688 (n=8-12). (**b**) Percentages of Foxp3^+^ Tregs (*top panel*) and geometric mean fluorescence intensity (gMFI) of Foxp3 (*bottom panel*). (**c)** Representative flow cytometry dot plots of RORγt expression among Foxp3^+^ Tregs in colon and small intestinal lamina propria. (**d)** Percentages of RORγt expression by Foxp3^+^ Tregs. (**e)** Absolute numbers of RORγt^+^Foxp3^+^ Tregs, RORγt^−^Foxp3^+^ Tregs, RORγt^+^Foxp3^−^ Th17 cells in colon lamina propria (cLP). Each scatter dot plot in bar graphs represents data from one mouse. Data shown as mean ± SD are pooled from four independent experiments and analysed by ANOVA with post-hoc Holm-Sidak’s multiple comparisons test (**b, d, e**). \**P*<0.05, \*\**P*<0.01, \*\*\**P*<0.001, \*\*\*\**P*<0.0001, and ns=not significant.

It has recently been reported that a subpopulation of intestinal Tregs that express the transcription factor retinoid-related orphan receptor gamma t (RORγt), namely RORγt^+^Foxp3^+^ Tregs, inhibit intestinal inflammation with greater suppressive potential than RORγt^−^Foxp3^+^ Tregs ^6,7,33^. We therefore examined the effects of PGE_2_ on RORγt^+^Foxp3^+^ Tregs and found that PGE_2_-EP4 signaling notably down-regulated RORγt^+^Foxp3^+^ Tregs in the colon, but not in mLN or spleen (**Fig. 2c, d)**. Furthermore, PGE_2_-EP4 signaling also decreased absolute numbers of RORγt^+^Foxp3^+^ Tregs, but not the numbers of RORγt^−^Foxp3^+^ Tregs or RORγt^+^Foxp3^−^ Th17 cells (**Fig. 2e**). These results indicate that PGE_2_-EP4 signaling suppresses Tregs with the greatest potency in the intestine.

### The gut microbiota is involved in PGE_2_ suppression of intestinal Tregs

To examine whether PGE_2_-EP4 signaling inhibits intestinal Tregs in the steady state via direct or indirect actions on T cells, we generated Lck-Cre driven EP4 conditional knockout mice by crossing EP4-flox mice to Lck-Cre mice (i.e. Lck^Cre^EP4^fl/fl^ mice) to delete EP4 expression in Lck-expressing T cells^29^. Lck^Cre^EP4^fl/fl^ and control mice had comparable colonic Treg accumulation in the steady state (**Fig. 3a)**, suggesting that PGE_2_ inhibits intestinal Treg accumulation in the steady state independent of EP4 signaling in T cells. The gut microbiota is crucial for the development of intestinal Tregs, especially the RORγt^+^Foxp3^+^ Treg subset ^6,7,33^. We therefore investigated whether the gut microbiota is involved in PGE_2_-dependent control of intestinal Tregs. We analyzed colonic Tregs from WT mice in which the gut microbiota had been depleted by antibiotics, or from MyD88/TRIF double knockout mice that were unable to sense microbial signals. While inhibition of endogenous PGE_2_ by indomethacin increased both the frequencies and numbers of colonic Tregs (especially the sub-population of RORγt^+^Foxp3^+^ Tregs) but not RORγt^+^Foxp3^−^ Th17 cells in WT mice that have been treated with vehicle control, there was no increased accumulation of colonic Tregs by indomethacin in antibiotic-treated WT mice nor in those mice with dual deficiency of MyD88 and TRIF (**Fig. 3b**). Similarly, activation of EP4 reduced indomethacin-dependent increase in colonic RORγt^+^Foxp3^+^ Tregs in vehicle-treated, rather than antibiotic-treated, mice (**Fig. 3b**). These results suggest the involvement of commensal microbiota in the PGE_2_-dependent control of intestinal Tregs.

**Figure 3.**
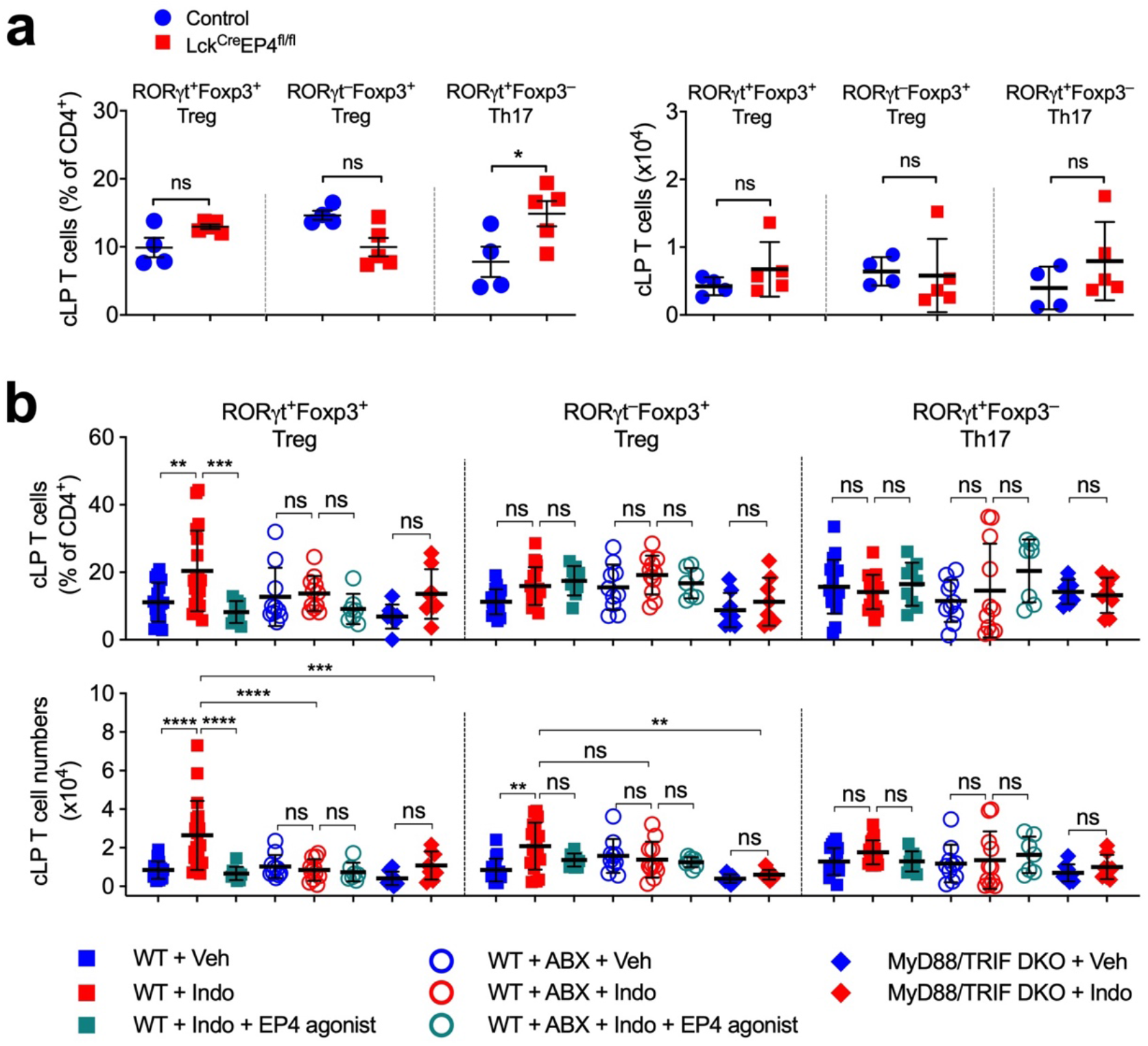
The gut microbiota is involved in PGE_2_-EP4 signaling regulation of intestinal Tregs. **(a)** Percentages and numbers of RORγt^+^Foxp3^+^ Tregs, RORγt^−^Foxp3^+^ Tregs, RORγt^+^Foxp3^−^ Th17 cells in colon lamina propria (cLP) of Lck^Cre^EP4^fl/fl^ (n=5) and control (n=4) mice determined by flow cytometry. **(b)** Percentages and numbers of RORγt^+^Foxp3^+^ Tregs, RORγt^−^Foxp3^+^ Tregs and RORγt^+^Foxp3^−^ Th17 cells in cLP of wild-type (WT) and MyD88/TRIF double knockout (DKO) mice that were treated with vehicle (Veh), indomethacin (Indo), or indomethacin plus an EP4 agonist, L-902,688 (n=7-18). Antibiotics (ABX) were administered to WT mice in drinking water for 2 weeks started 1 week before receiving indomethacin or vehicle. Each scatter dot plot in bar graphs represents data from one mouse. Data shown as mean ± SD are pooled from two (**a**) or five (**b**) experiments and analysed by ANOVA with post-hoc Holm-Sidak’s multiple comparisons test. \**P*<0.05, \*\**P*<0.01, \*\*\**P*<0.001, \*\*\*\**P*<0.0001, and ns=not significant.

### PGE_2_–EP4 signaling changes specific gut microbial populations

Use of NSAIDs has been reported to induce changes in the gut microbiota composition in humans and rodents ^34–36^. To examine whether PGE_2_-EP4 signaling modulates the gut microbiota, we collected cecal contents from mice that had been treated with indomethacin or that had been co-treated with indomethacin and an EP4 agonist, and performed 16S rRNA gene metabarcoding to study microbiota composition. Principal component analysis (PCoA) suggested that there were no differences in overall β-diversity of gut microbiota signatures measured by unweighted UniFrac distances among the three groups (**Fig. 4a**). The analysis of α-diversity indices (i.e., richness and evenness) showed that the three groups had also comparable observed OTUs, Chao1 index, Shannon diversity and Inverse Simpson indexes (**Fig. 4b**). But there were trends showing that the indomethacin+EP4 agonist-treated group had higher observed OTUs and Chao1 index (**Fig. 4b**). We then asked whether PGE_2_-EP4 signaling alters specific bacterial communities. We found that treatment with indomethacin increased the abundance of the *Firmicutes* phylum and reduced the abundance of the *Bacteroidetes* phylum, and the changes in phylum-level microbiota composition by indomethacin was slightly reversed by co-treatment with the EP4 agonist (**Fig. 4c**). Mice treated with indomethacin increased some aggressive species like the *Rikenella* and *Escherichia* genus, which was reduced by EP4 agonist (**Fig. 4d**). This may be associated with the fact that NSAIDs potentially damage the gut epithelium and PGE_2_-EP4 signaling protects against the damage. On the other side, we also found that indomethacin increased, but EP4 agonist markedly reduced, several SCFA-producing bacteria belonging to the *Muribaculaceae* family or the *Clostrium* cluster XIVa (e.g., *Lachnospiraceae, Ruminococcaceae*)^5^ (**Fig. 4d**). Furthermore, *Anaeroplasma bactoclasticum*, which promotes expression of immune-regulatory TGF-β in the gut ^37^, was also upregulated by indomethacin and reduced by the EP4 agonist (**Fig. 4d**). SCFA-producing bacteria like *Clostria* play critical roles in intestinal Treg induction and accumulation. To validate the 16S rRNA gene sequencing results, we employed real-time quantitative polymerase chain reaction (qPCR) to detect gene expression of SCFA-producing bacteria in mice from independent cohorts. As confirmed, activation of EP4 significantly reduced the phylum *Firmicutes* and increased the phylum *Bacteroidetes* (**Fig. 4e**). In agreement with 16S RNA sequencing results, EP4 activation notably decreased the amounts of *Clostridia* including total *Clostridium XIVa, Clostridium sp*., *C. coccoides* and *ASF500* (*Ruminococcaceae*) and *ASF360* (*Lactobacillus sp*.) (**Fig. 4e**). Taken together, our results suggest that PGE_2_-EP4 signaling alters SCFA-producing microbiota, which may result in reduction of intestinal Treg accumulation.

**Figure 4.**
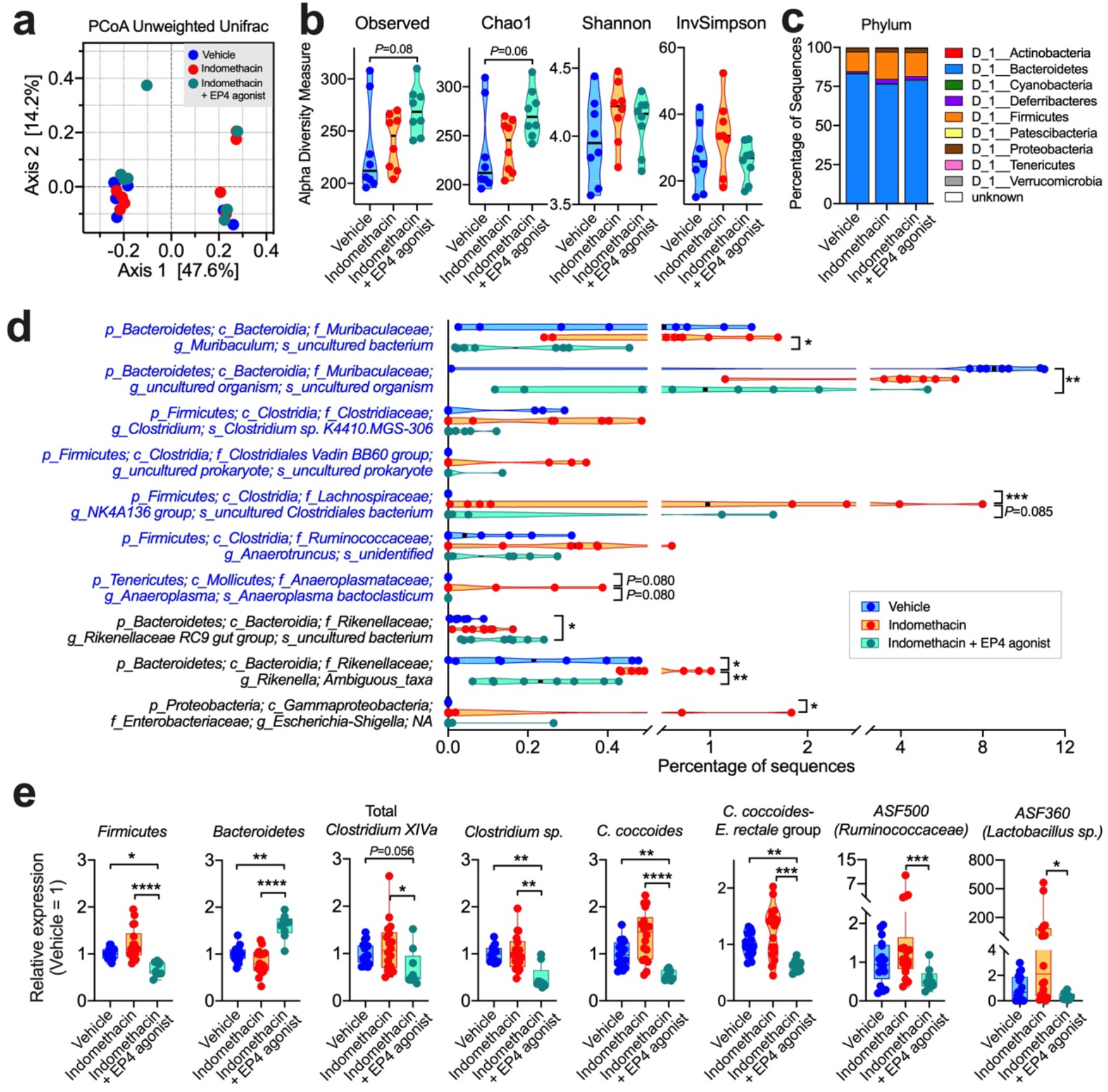
PGE_2_–EP4 signaling alters the gut microbiota. **(a-d)** DNA extracts of the cecal contents from C57BL/6 mice receiving vehicle control or indomethacin plus EP4 agonist for 5 days (n=8 per group) were analyzed by 16S rRNA gene sequencing. (**a**) Principal component analysis (PCoA) of cecal microbiota structure as measured by unweighted UniFrac distances among the three groups. (**b)** Alpha-diversity of the gut microbiome as measured by observed OTUs, Chao1 index, Shannon diversity and Inverse Simpson (InvSimpson) index. (**c)** Phylum-level microbiota composition expressed as relative abundances. (**d)** Bacterial species changed in relative abundance among different groups. (**e)** 16S rRNA gene expression of indicated commensal microbiota in cecal contents obtained from C57BL/6 mice treated with vehicle (n=17), indomethacin (n=18) or indomethacin plus the EP4 agonist L-902,688 (n=9) for 5 days. Relative abundance of each group of bacteria to total cecal bacteria was determined by quantitative real-time polymerase chain reaction (qPCR) and normalised to the vehicle group. Each scatter dot plot represents data from one mouse (**a, b, d, e**). Data plotted in box and whiskers bar graphs are pooled from five experiments (**e**) and analysed by non-parametric Kruskal-Wallis test with post-hoc Dunn’s multiple comparisons test. \**P*<0.05, \*\**P*<0.01, \*\*\**P*<0.001, \*\*\*\**P*<0.0001, and ns=not significant.

### PGE_2_–EP4 signaling-modified gut microbiota suppresses mucosal Tregs and promotes intestinal inflammation

To investigate whether PGE_2_-modified microbiota modulates intestinal Treg responses and inflammation, we adoptively transferred cecal microbiota obtained from WT C57BL/6 mice that had been treated with indomethacin, or indomethacin plus an EP4 agonist, or vehicle control into recipient WT C57BL/6 mice (**Fig. 5a**). Recipient mice were pre-treated with antibiotics in drinking water for 2 weeks before receiving transplantation of microbiota, and then received normal drinking water for another 10 days followed by euthanizing to analyze colonic immune cell responses at the steady state. Some recipient mice received normal drinking water after stop of antibiotic-treatment for another 8 days followed by receiving dextran sulfate sodium (DSS) in drinking water for further 6 days (**Fig. 5a**). Compared to mice that had received cecal microbiota from vehicle-treated mice, mice transplanted with cecal microbiota from indomethacin-treated mice had increased Tregs as well as Foxp3 expression at single cell levels in the steady state (**Fig. 5b**). This was associated with prevented body weight loss, reduced disease activity index, and increased colon length under DSS-induced inflammatory conditions in mice that had received cecal microbiota from indomethacin-treated mice (**Fig. 5c, d**). In contrast, mice transplanted with cecal microbiota from mice that have been pre-treated with both indomethacin and the EP4 agonist had a trend to reduce colon Tregs and Foxp3 expression compared to mice received cecal microbiota from mice that have been pre-treated with indomethacin (**Fig. 5b**). Furthermore, the severity of colitis in mice transplanted with cecal microbiota from mice that have been pre-treated with both indomethacin and EP4 agonist was similar to mice that had received cecal microbiota from vehicle-treated mice, but was significantly greater than that in mice received cecal microbiota from indomethacin-treated mice (**Fig. 5c-e**). Histological analysis showed near normal proximal and distal colonic mucosa, or only scattered mild inflammatory changes, in mice transplanted with cecal microbiota from indomethacin-treated mice **(Fig. 5f)**. There was widespread and variably severe mucosal ulceration with patches of almost complete loss of the crypt epithelium, with marked infiltration of fibrotic mucosal tissue by both acute and chronic inflammatory cells in mice that had received cecal microbiota from vehicle-treated mice, with more severe inflammatory and ulcerative changes in the distal colon compared with the proximal colon. In contrast, there was a less severe pattern of inflammation with more variable mild to moderate inflammatory cell infiltration with only patchy partial loss of mucosal crypt epithelium in those mice transplanted with cecal microbiota from indomethacin- and EP4 agonist-treated mice, again with greater inflammatory changes in the distal colon compared with the proximal colon (**Fig. 5f**). This was associated with reduced T cell infiltration to colon LP in mice that have received cecal microbiota from indomethacin-treated mice compared to the other two groups (**Fig. 5g**). Together, these results suggest that PGE_2_-EP4 signaling-mediated modification of the gut microbiota contributes to inhibition of mucosal Tregs and exacerbation of intestinal inflammation.

**Figure 5.**
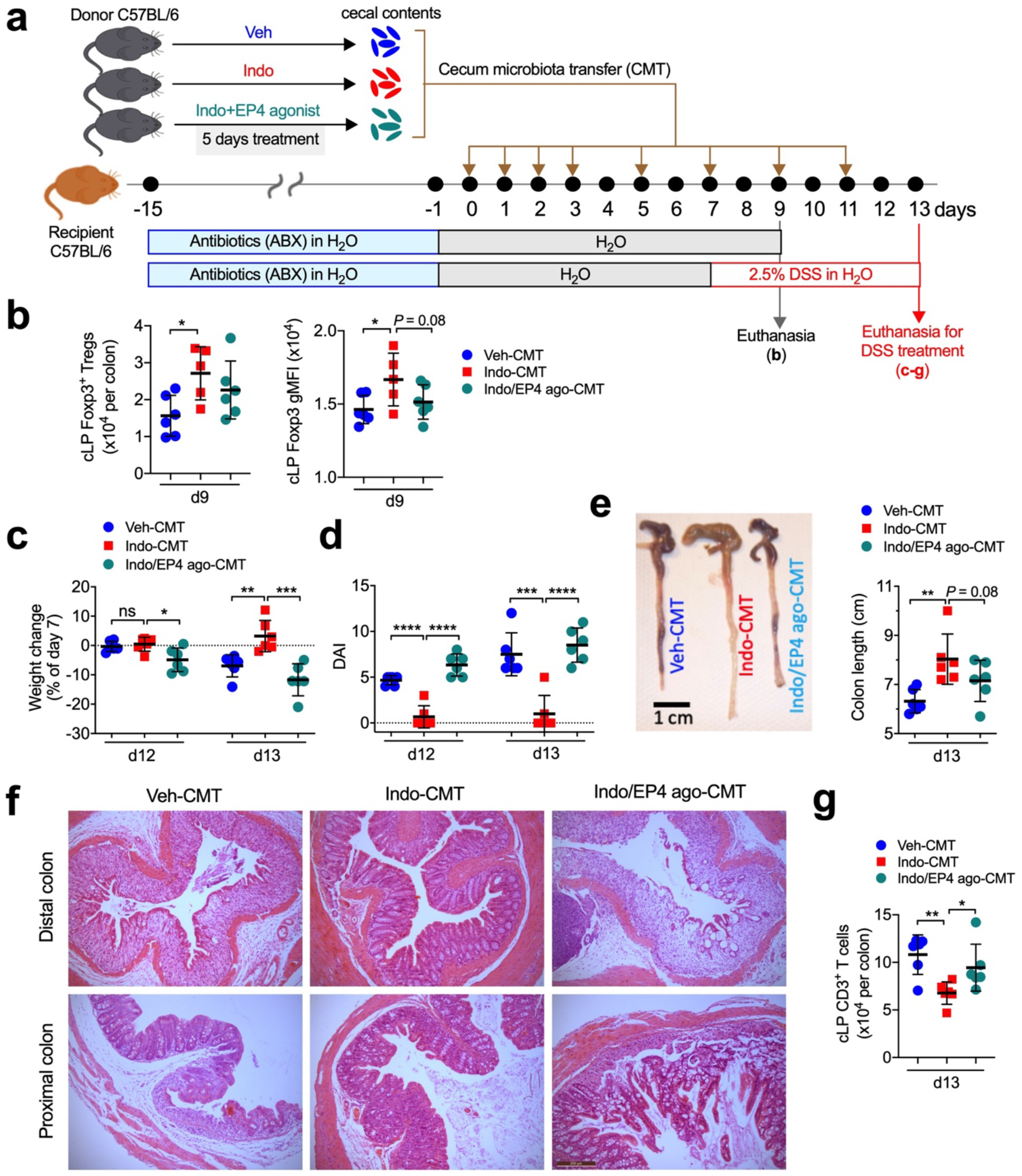
Altered gut microbiota by PGE_2_-EP4 signaling suppresses intestinal Treg responses and promotes intestinal inflammation. **(a)** Experimental schematic diagram of cecal microbiota transplantation (CMT). Recipient C57BL/6 mice were pre-treated with antibiotics for 2 weeks and rest for 1 day before receiving fresh cecal microbiota collected from C57BL/6 mice that had been treated with vehicle (Veh-CMT), indomethacin (Indo-CMT), or indomethacin plus EP4 agonist, L-902,688 (Indo/EP4 ago-CMT) for 5 days. After antibiotics treatment, some recipient mice were given back normal drinking water for 9 days before euthanised for analysis of colon immune cells (for **b**), while some other recipient mice were given back normal drinking water for 7 days followed by administration with dextran sulfate sodium (DSS) in drinking water to induce colonic inflammation (for **c-f)**. (**b)** Numbers of colon lamina propria (cLP) Foxp3^+^ Tregs (left) and Foxp3 geometric mean fluorescence intensity (gMFI, right) in the recipient mice on day 9 post CMT (n=5-6 each group). (**c)** Changes in body weight of recipient mice showing percentages of that at the beginning of DSS treatment (i.e. day 7). (**d)** Disease activity index (DAI) of the recipient mice (right) (n=6 each group). (**e)** Representative images of the cecum and colon tissues (left) and colon lengths (right) of the recipient mice on day 13. (**f)** Representative histological (H&E stained) images of distal (upper row) and proximal (lower row) colons of the recipient mice (all at same magnification 100x). Bar scale, 200 μm. (**g)** Numbers of infiltrated CD3^+^ T cells in colon LP of the recipient mice. Each scatter dot plot in bar graphs represents data from one mouse. Data shown as mean ± SD are analysed by ANOVA with post-hoc Holm-Sidak’s multiple comparisons test (**b-e,g**). \**P*<0.05, \*\**P*<0.01, \*\*\**P*<0.001, \*\*\*\**P*<0.0001, and ns=not significant.

### PGE_2_ regulates intestinal MNPs required for the control of Tregs

MNPs are critical to mediate the microbiota-dependent generation of intestinal Foxp3^+^ Tregs 38-40. To further study the interplay between PGE_2_ and the gut microbiota in the control of intestinal Tregs, we examined MNPs in the colon. Inhibition of endogenous PGE_2_ increased both the frequency and number of colonic CD11c^+^MHC II^+^CD11b^+^ MNPs in WT mice that have been treated with vehicle control, and this was again prevented by co-administration of the EP4 agonist (**Fig. 6a**). CD11c^+^MHC II^+^CD11b^−^ MNPs were not affected by either indomethacin or EP4 activation (**Fig. 6a**). The effects of indomethacin and the EP4 agonist on colonic CD11c^+^MHC II^+^CD11b^+^ MNPs were invisible in colons of antibiotic-treated mice or in MyD88/TRIF-double deficient mice (**Fig. 6a**). Moreover, transfer of gut microbiota from mice that had been treated with indomethacin increased colonic CD11c^+^MHC II^+^CD11b^+^ MNPs in host mice, and this was again reduced by transfer of gut microbiota from mice that had been treated with both indomethacin and the EP4 agonist (**Fig. 6b**). These results indicate that PGE_2_-EP4 signaling suppresses intestinal MNPs through modulating the gut microbiota.

**Figure 6.**
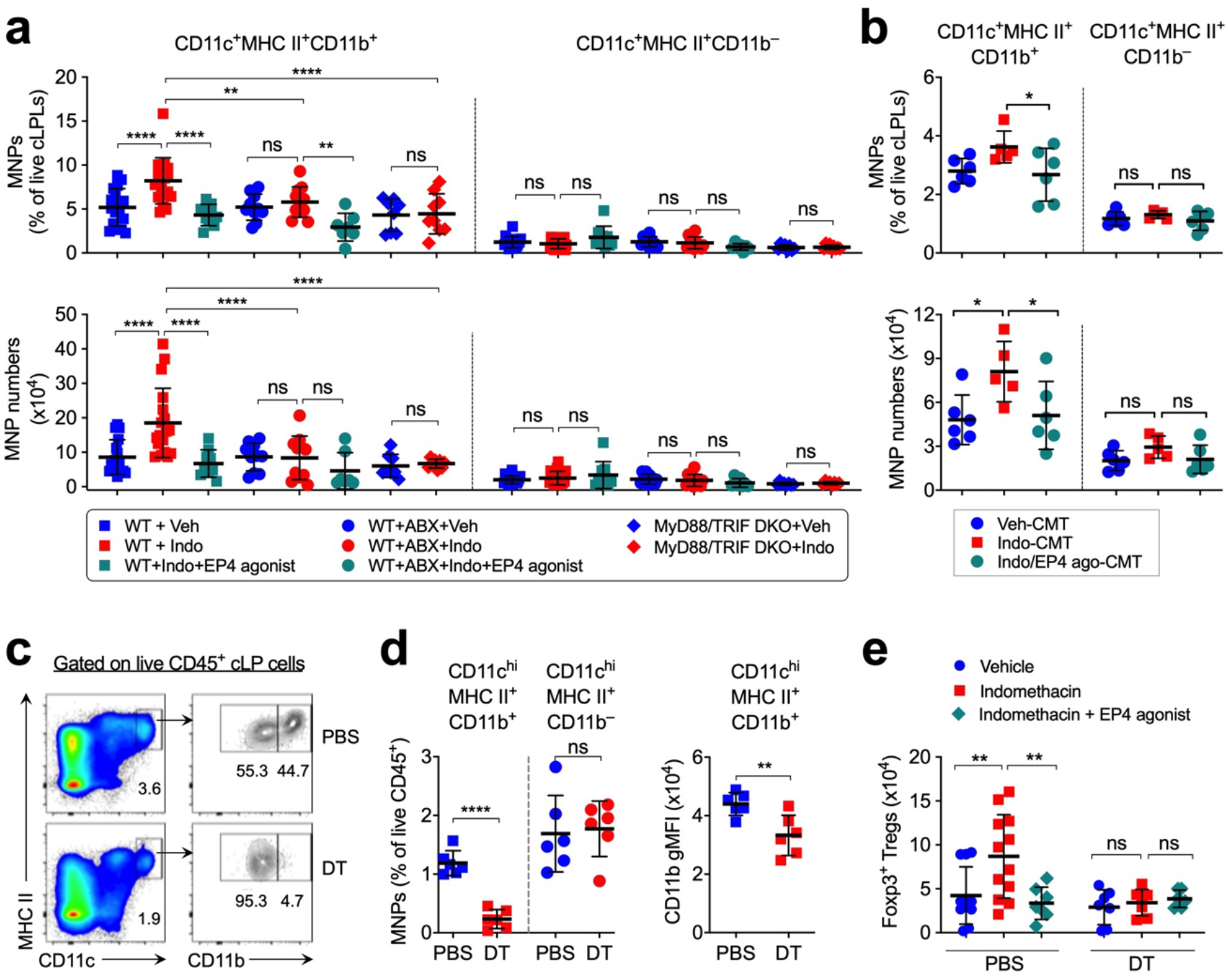
PGE_2_ inhibits intestinal mononuclear phagocytes (MNPs) via the gut microbiota, required for control of Tregs. **(a)** Percentages (upper) and numbers (lower) of colon lamina propria (cLP) CD11c^+^MHC II^+^CD11b^+^ and CD11c^+^MHC II^+^CD11b^−^ MNPs from C57BL/6 wild-type (WT) and MyD88/TRIF double knockout (DKO) mice treated with vehicle (Veh), indomethacin (Indo), or indomethacin plus an EP4 agonist L-902,688 (n=7-18). Antibiotics (ABX) were administered to WT mice in drinking water for 2 weeks started 1 week before receiving indomethacin or vehicle. (**b)** Percentages (upper) and numbers (lower) of colon LP CD11c^+^MHC II^+^CD11b^+^ and CD11c^+^MHC II^+^CD11b^−^ MNPs in mice that were treated with antibiotics for 2 weeks followed by transfer of cecal microbiota harvested from mice that had been administered with vehicle (Veh-CMT), indomethacin (Indo-CMT), or indomethacin plus EP4 agonist (Indo/EP4 ago-CMT) (n=5-6). **(c,d)** Representative flow cytometry dot-plots (**c**) and percentages (**d**) of colon LP CD11c^+^MHC II^+^CD11b^+^ and CD11c^+^MHC II^+^CD11b^−^ MNPs at colon lamina propria of CD11b-DTR mice administered with diphtheria toxin (DT) or PBS (n=6 each group). CD11b geometric mean fluorescence intensity (gMFI) of CD11c^+^MHC II^+^CD11b^+^ MNPs are also shown (**d**, right). (**e)** Numbers of colon LP CD45^+^CD3^+^CD4^+^Foxp3^+^ Tregs in CD11b-DTR mice treated with DT or PBS together with vehicle control, indomethacin, or indomethacin plus EP4 agonist (n=6-12). Each scatter dot plot in bar graphs represents data from one mouse. Data shown as mean ± SD are pooled from five (**a**), one (**b**) or three (**d,e**) independent experiments and analysed by ANOVA with post-hoc Holm-Sidak’s multiple comparisons test (**a, b, e**) or two-tailed unpaired student *t*-test (**d**). \**P*<0.05, \*\**P*<0.01, \*\*\**P*<0.001, \*\*\*\**P*<0.0001, and ns=not significant.

Next, we asked whether CD11c^+^MHC II^+^CD11b^+^ MNPs mediate PGE_2_ suppression of intestinal Tregs. To address this, we took advantage of mice that have CD11b promoter-driven expression of the human diphtheria toxin (DT) receptor (CD11b-DTR mice). Injection of DT selectively deplete colonic LP CD11c^+^MHC II^+^CD11b^+^ MNPs (by ~90%), rather than CD11c^+^MHC II^+^CD11b^−^ MNPs, as well as markedly reduced CD11b expression at the single cell level in colonic CD11c^+^MHC II^+^CD11b^+^ MNPs (**Fig. 6c,d**). While indomethacin increased and EP4 agonist decreased colonic Foxp3^+^ Tregs in PBS-treated CD11b-DTR mice, respectively, neither indomethacin nor EP4 agonist affected colonic Tregs in DT-treated CD11b-DTR mice (**Fig. 6e**). These results suggest that CD11c^+^MHC II^+^CD11b^+^ MNPs mediate PGE_2_ suppression of intestinal Tregs. Furthermore, the C-C chemokine receptor type 2 (CCR2) mediates migration of monocyte-derived macrophages to the intestine. Mice deficient in CCR2 had reduced intestinal CD11c^+^MHC II^+^CD11b^+^ MNP numbers (by ~50%) and CD11b expression at the single cell level compared to WT mice (**Supplementary Fig. 3a**). However, CCR2 deficiency did not prevent, if not enhanced, indomethacin-dependent increase in colonic Foxp3^+^ Tregs (**Supplementary Fig. 3b**). These results indicate that PGE_2_-dependent regulation of intestinal Treg responses is likely to be mediated by gut resident but not monocyte-derived migratory MNPs.

### Type I interferon signaling mediates PGE_2_-dependent suppression of intestinal MNPs and Tregs

MNPs mediate intestinal Treg development and expansion by producing soluble mediators such as type I IFNs ^41,42^. To test whether PGE_2_ regulates type I IFN signaling in intestinal CD11c^+^MHC II^+^CD11b^+^ MNPs, we cultured bone morrow-derived dendritic cells (BMDCs) without (medium only) or with cecal microbial products (CMPs, which might include pathogen- or microbe-associated molecular patterns including gut microbial metabolites) obtained from mice that had been treated with vehicle control, indomethacin, or indomethacin plus an EP4 agonist. We then measured mRNA expression of type I IFN signaling pathway genes in BMDCs by real-time qPCR. BMDCs cultured with CMPs from control mice were characterized by higher expression of type I IFN (e.g. *Ifnb*) and the downstream genes (e.g. *Irf7* and *Isg15*) compared to that cultured with medium only (**Fig. 7a**). Expression of *Irf7* and *Isg15* was further increased in BMDCs cultured with CMPs obtained from mice that had been treated with indomethacin compared to that cultured with CMPs obtained from control mice (**Fig. 7a**). In contrast, mRNA expression levels of all these genes were markedly down-regulated in BMDCs that had been cultured with CMPs obtained from mice that have been co-treated with indomethacin and EP4 agonist (**Fig. 7a**). Moreover, colonic CD11c^+^MHC II^+^ MNPs sorted from indomethacin-treated mice expressed higher levels of type I IFN signaling pathway genes (e.g. *Ifnb*, *Irf7* and *Isg15*) compared to MNPs sorted from colons of vehicle-treated mice (**Fig. 7b**). Colonic MNPs in mice treated with indomethacin had higher levels of phosphorylated STAT1 than that in vehicle-treated mice **(Fig. 7c)**. These results suggest that modulation of gut microbial products by PGE_2_-EP4 signaling inhibits type I IFN production and signaling in MNPs. Similarly, colonic Tregs from indomethacin-treated mice also had more Tregs expressing phosphorylated STAT1 compared to vehicle-treated mice (**Fig. 7d**). We next examined whether type I IFN signaling is required for PGE_2_ control of intestinal Tregs using IFN-α/β receptor α chain (*Ifnar*) deficient mice. Again, indomethacin increased colonic CD11c^+^MHC II^+^CD11b^+^ MNPs and RORγt^+^Foxp3^+^ Tregs, and this was prevented by EP4 agonist in WT mice (**Fig. 7e, f**). However, neither indomethacin nor EP4 agonist had effects on colonic CD11c^+^MHC II^+^CD11b^+^ MNPs and RORγt^+^Foxp3^+^ Tregs in *Ifnar*-deficient mice (**Fig. 7e, f**). These results thus suggest that the PGE_2_-modified microbiota suppresses intestinal Tregs by down-regulating type I IFN signaling via MNPs.

**Figure 7.**
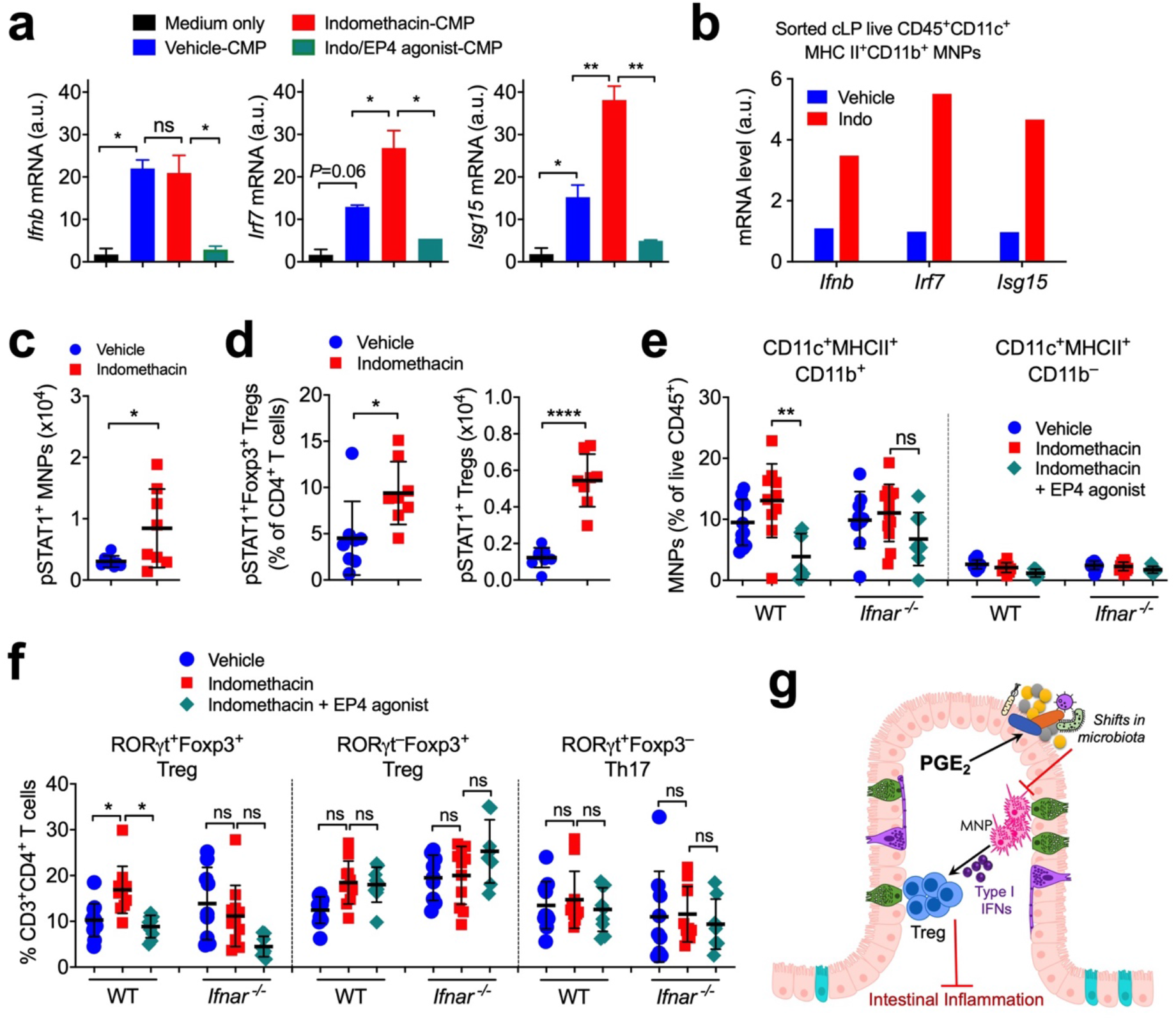
Type I IFNs produced by mononuclear phagocytes (MNPs) mediate PGE_2_-dependent suppression of intestinal Tregs. **(a)** Gene expression of *Ifnb*, *Irf7* and *Isg15* in bone marrow-derived dendritic cells (BMDCs) cultured for 6 h with cecal microbial products (CMPs) that were isolated from mice treated with vehicle (Veh-CMP), indomethacin (Indo-CMP), or indomethacin plus EP4 agonist, L902,688 (Indo/EP4 ago-CMP) for 5 days. Real-time qPCR results were normalized to *Gapdh* and relative to the group that BMDCs were cultured with medium only. Data shown as mean ± SEM of duplicates are representative of two independent experiments and analysed by ANOVA with post-hoc Holm-Sidak’s multiple comparisons test. \**P*<0.05, \*\**P*<0.01, and ns=not significant. (**b)** Gene expression of *Ifnb*, *Irf7* and *Isg15* in CD45^+^CD3^−^B220^−^CD11c^+^MHC II^+^ MNPs sorted from colon lamina propria (cLP) of mice treated with vehicle (Veh) or indomethacin (Indo) for 5 days. Real-time qPCR results were normalized to *Gapdh* and relative to MNPs isolated from vehicle-treated mice. Each sample was pooled from 3-4 mice. Data shown as mean gene expression of technical du- or triplicates are representative of two independent experiments. (**c)** Numbers of pSTAT1^+^ MNPs in the colon from mice treated with vehicle or indomethacin (n=8 each) for 5 days. (**d)** Percentages and numbers of cLP pSTAT1^+^Foxp3^+^ Tregs. Each scatter dot plot represents data from one mouse. Data shown as mean ± SD are pooled from two independent experiments and analysed by two-tailed unpaired student *t*-test. \**P*<0.05 and \*\*\*\**P*<0.0001. (**e)** Percentages of cLP CD11c^+^MHC II^+^CD11b^+^ and CD11c^+^MHC II^+^CD11b^−^ MNP subsets in wild-type (WT) and *Ifnar* knockout (*Ifnar^−/−^*) mice treated with vehicle, indomethacin, or indomethacin plus EP4 agonist, L-902,688 for 5 days (n=7-12). (**f)** Percentages of cLP RORγt^+^Foxp3^+^ Tregs, RORγt^−^Foxp3^+^ Tregs and RORγt^+^Foxp3^−^ Th17 cells. Each scatter dot plot represents data from one mouse. Data shown as mean ± SD are pooled from four independent experiments and analysed by ANOVA with post-hoc Holm-Sidak’s multiple comparisons test. \**P*<0.05, \*\**P*<0.01, and ns=not significant. (**g)** Diagram illustrating how PGE_2_ suppresses intestinal Treg responses through shaping the gut microbiota composition and modulation of MNP functions.

## Discussion

In this report, we have shown that the well-known lipid mediator of inflammation, PGE_2_, acts as a pivotal controller of intestinal Treg responses to foster intestinal inflammation. Endogenous PGE_2_-EP4 signaling alters the gut microbial community, e.g. reducing SCFA-producing bacteria, which in turn down-regulates MNP production of type I IFNs, leading to repression of intestinal Treg accumulation and augmentation of intestinal inflammation (**Fig. 7g**). In contrast, blockade of PGE_2_ signaling (e.g. by NSAIDs) increases beneficial microbes and augments MNP production of IFNs, resulting in expansion of mucosal Trges and limitation of intestinal inflammation.

Blockade of PGE_2_ biosynthesis by NSAIDs alters the gut microbiota composition by expanding the beneficial SCFA-producing microbes. Transfer of the gut microbiota from NSAID-treated mice efficiently prevented intestinal inflammation associated with increased intestinal Tregs. This effect of NSAIDs can be counteracted by activation of PGE_2_ signaling through EP4. Our results showing that PGE_2_-EP4 signaling reduced *Firmicutes* but increased *Bacteroidetes* at the phylum level may have implications in modulation of intestinal inflammation as an increased ratio of *Firmicutes* to *Bacteroidetes* has been reported to be related to anti-inflammatory potential under various inflammatory conditions ^3^. Furthermore, our findings on the manipulation of the gut microbiota in mice are in keeping with observations in humans showing that ingestion of antipyretic analgesics (e.g. PG-inhibiting NSAIDs or paracetamol) increased the abundance of beneficial gut commensals such as *Verrucomicrobia* and SCFA-producing bacteria like *Butyrivibrio* and *Clostridiaceae* ^36^. A recent report performing *in vitro* culture of commensal bacteria suggested that most NSAIDs, including indomethacin and aspirin, were unlikely to impact bacterial growth on various microbial strains^43^. Given that PGE_2_ may promote growth of some bacteria such as *Escherichia coli* ^44^, further studies are needed to decipher whether PGE_2_ acts directly on the gut microbiota or indirectly on host cells to regulate growth or survival of specific gut commensal bacterial strains.

Mice with EP4 deficiency in T cells had comparable numbers of intestinal Tregs to EP4-sufficient mice, indicating that the inhibitory effect of PGE_2_ on the accumulation of intestinal Tregs was not mediated by EP4 signaling in T cells. Instead, MNPs seem to mediate PGE_2_ suppression of intestinal Tregs. Indeed, alteration of PGE_2_-EP4 signaling (e.g. using COX inhibitors or EP4 agonist) changes the numbers of intestinal CD11c^+^MHC II^+^CD11b^+^ MNPs, and depletion of CD11b^+^ MNPs prevented PGE_2_-dependent regulation of intestinal Tregs. While CCR2-expressing monocyte-derived MNPs are critical for intestinal Th17 cell responses 45, these migratory MNPs are unlikely to be involved in PGE_2_-dependent regulation of intestinal Treg responses, indicating a role for gut resident MNPs. Furthermore, PGE_2_-modified gut microbiota controls MNP production of type I IFNs, which in turn mediates intestinal Tregs and MNPs themselves. This is consistent with a recent report showing that gut commensals stimulated CD11b^+^ dendritic cells to produce interferon-β, which augmented the proliferation of intestinal Tregs ^41^.

Over-activation of the PGE_2_ pathway including PGE_2_ biosynthesis and its receptor signaling is an outstanding marker in most, if not all, human inflammatory conditions such as multiple sclerosis, rheumatoid arthritis, IBD and cancers^14^. These results not only imply PGE_2_ as a common inflammatory mediator for immune inflammation by balancing T cell responses, i.e. enhancing Treg but limiting effector T cell responses^28,29^, but also suggest that targeting the PGE_2_ pathway may potentially be of beneficial in the control of intestinal inflammation. Our findings are further supported by a recent study demonstrating that PGE_2_ exacerbates TNF-induced inflammatory responses in human intestinal epithelial cells from patients with IBD who are resistant to TNF inhibitor therapy ^46^. Moreover, recent studies including ourselves have suggested that lack of PGE_2_-EP4 signaling in T cells reduced both chemical-triggered acute and naïve T cell transfer-induced chronic intestinal inflammation, associated with reduction of inflammatory Th1 and/or Th17 cell responses ^29,47,48^. This work thus advances our understanding that PGE_2_ mediates intestinal inflammation through modulating the network of the gut microbiota and the host immune system involving both innate MNPs and adaptive T cell responses.

Collectively, we have updated our understanding of the roles for PGE_2_ in the settings of intestinal inflammation and proposed the immune mechanisms underpinning its deleterious function. Our results provide an explanation for human genetic findings of the association between EP4 gene polymorphisms and IBD susceptibility. This work suggests that targeting the PGE_2_-EP4-microbiota-MNP-Treg cascade might be a promising therapeutic strategy for intestinal inflammation.

## Methods

### Mice

*Rag1^−/−^*, MyD88^−/−^TRIF^−/−^, Ifnar^−/− 49^, CD11b-DTR, Ccr2^−/−^, Lck^Cre^EP4^fl/fl 29^, and wild-type C57BL/6 mice were bred and maintained under specific pathogen-free conditions in accredited animal facilities at the University of Edinburgh. Wild-type mice were purchased from Harlan UK or bred in our own animal facilities. Age-(>7-weeks old) and sex-matched mice were used. Mice were randomly allocated into different groups and analysed individually. No mice were excluded from the analysis except exclusions due to technical errors in preparation of intestinal lamina propria leukocytes. All experiments were conducted in accordance with the UK Animals (Scientific Procedures) Act of 1986 with local ethical approval from the University of Edinburgh Animal Welfare and Ethical Review Body (AWERB).

### Cecal microbiota transplantation (CMT), colitis and treatments with small molecular compounds and antibiotics

Recipient SPF WT C57BL/6 mice were pre-treated with antibiotics for 2 weeks and rest for 1 day before receiving fresh cecal microbiota collected from SPF WT C57BL/6 mice that had been treated with vehicle, indomethacin (5 mg/kg/day in drinking water), or indomethacin plus EP4 agonist L-902,688 (10 μg/mouse/d via daily intraperitoneal injection) for 5 days. After antibiotics treatment, some recipient mice were given normal drinking water for another 9 days before euthanised for analysis of colon immune cells in the steady state, while other recipient mice were administered with normal drinking water for 7 days followed by 2.5% (w/v) of dextran sulfate sodium (DSS, MW 36-50kDa, MP Biochemical) in drinking water for 6 consecutive days to induce colonic inflammation. The disease activity index (DAI) was scored by the following system. Body weight: 0 (no or <1% weight loss compared to d 0 body weight), 1 (1-5% weight loss), 2 (5-10% weight loss), 3 (10-20% weight loss), and 4 (>20% weight loss); bleeding: 0 (no bleeding), 1 (blood present in/on faeces), 2 (visible blood in rectum), and 4 (visible blood on fur); stool consistency: 0 (well-formed/normal stool), 1 (pasty/semi-formed stool), 2 (pasty stool with some blood), 3 (diarrhea that does not adhere to anus), and 4 (diarhoea that does adhere to anus); and general appearance: 0 (normal), 1 (piloerection only), 2 (piloerection and lethargy), 4 (atoxic, motionless and sunken eyes). Mice were immediately culled when body weight loss was greater that 25% or the total colitis score is 12 or higher. Indomethacin (5 mg/kg/d or indicated doses) and vehicle (0.5% EtOH) were administrated through drinking water which was refreshed every 2-3 days. Butaprost (10 μg/injection, Abcam), L-902,688 (10 μg/injection, Cayman) and control (0.5% EtOH in PBS) was used by daily intraperitoneal injections. Antibiotics (containing ampicillin, gentamycin, metronidazole, neomycin, vancomycin, each 0.5 mg/ml) plus sucralose (4 mg/ml) was used in drinking water. Diphtheria toxin (DT, 25 ng/g bogy weight, Sigma) was injected intraperitoneally every 48 hours.

### Oxylipin analysis

Small intestine and colon samples were weighed and homogenized with ceramic beads in 1ml anti-oxidation buffer containing 100μM diethylenetriaminepentaacetic acid (DTPA) and 100μM M butylated hydroxytoluene (BHT) in phosphate buffered saline using a Bead Ruptor Elite for 2 × 30 second intervals at 6 m/s, under cooled nitrogen gas (4°C). Samples were spiked with 2.1-2.9ng of PGE_2_-d4, PGD_2_-d4, PGF_2_α-d4, TXB_2_-d4 standards (Cayman Chemical) prior to homogenization. Lipids were extracted by adding a 1.25 ml solvent mixture (1 M acetic acid/isopropanol/hexane; 2:20:30, v/v/v) to 0.5 ml supernatants in a glass extraction vial and vortexed for 30 sec. 1.25ml hexane was added to samples and after vortexing for 30 seconds, tubes were centrifuged (500 g for 5 min at 4 °C) to recover lipids in the upper hexane layer (aqueous phase), which was transferred to a clean tube. Aqueous samples were re-extracted as above by addition of 1.25 ml hexane, and upper layers were combined. Lipid extraction from the lower aqueous layer was then completed according to the Bligh and Dyer technique using sequential additions of methanol, chloroform and water, and the lower layer was recovered following centrifugation as above and combined with the upper layers from the first stage of extraction. Solvent was dried under vacuum and lipid extract was reconstituted in 200 μl HPLC grade methanol. Lipids were separated by liquid chromatography (LC) using a gradient of 30-100% B over 20 minutes (A: Water:Mob B 95:5 + 0.1% Acetic Acid, B: Acetonitrile: Methanol − 80:15 + 0.1% Acetic Acid) on an Eclipse Plus C18 Column (Agilent), and analysed on a Sciex QTRAP^®^ 6500 LC-MS/MS system. Source conditions: TEM 475°C, IS −4500, GS1 60, GS2 60, CUR 35. Lipid were detecting using MRM monitoring with the following parent to daughter ion transitions: PGD_1_ and PGE_1_ [M-H]- 353.2/317.2, PGD_2_, 8-iso PGE_2_ and PGE_2_ [M-H]- 351.2/271.1, PGF_2α_ [M-H]- 353.2/309.2, 6-keto PGF_1α_ [M-H]- 369.2/163.1, TXB_2_ [M-H]- 369.2/169.1, 13,14-dihydro-15-keto-PGE_2_ [M-H]- 351.2/235.1. Deuterated internal standards were monitored using parent to daughter ions transitions of: TXB_2_-d4 [M-H]- 373.2/173.1, PGE_2_-d4 and PGD_2_-d4 [M-H]- 355.2/275.1, PGF_2α_-d4 [-H]- 357.5/313.2. Chromatographic peaks were integrated using Multiquant 3.0.2 software (Sciex). Peaks were only selected when their intensity exceeded 3 times above the baseline noise. The ratio of analyte peak areas to internal standard was taken and lipids quantified using a standard curve made up and run at the same time as the samples. Each oxylipin was then standardized per mg of colon tissue.

### Histology

Intestine samples were fixed with 10% neutral buffered formalin solution (Sigma), embedded in paraffin, and 5 μm sections were used for staining with hematoxylin and eosin (H&E).

### Isolation and surface and intracellular staining of intestinal lamina propria leukocytes

Intestinal lamina propria cells were isolated as described previously ^19^. For surface staining, cells were first stained with the Fixable Viability Dye eFluor^®^ 780 (eBioscience) on ice for 30 min. After wash, cells were stained on ice for another 30 min with anti-CD45 (clone 30-F11), anti-CD3e (Clone 145-2C11), anti-CD4 (Clone GK1.5), anti-CD25 (clone PC61.5), anti-CD11c (clone N418), anti-CD11b (clone M1/70), anti-B220 (clone RA3-6B2), anti-mouse I-A/I-E Antibody (clone M5/114.15.2). For staining of transcription factors, cells were fixed in the Foxp3/Transcription Factor Fix Buffer (eBioscience) for 2 h or overnight followed by staining with anti-mouse Foxp3 (clone FJK-16s), anti-Mouse ROR-γt (clone B2D, eBioscience) and mouse anti-Stat1 (pY701) (clone 4a, BD UK) for at least 1 hour. All Abs were purchased from eBioscience, Biolegends or BD Bioscience. Flow cytometry was performed on the BD LSR Fortessa (BD Bioscience) and analyzed by FlowJo software (Tree Star).

### BMDC differentiation and stimulation

Bone marrow cells from femurs were cultured at 4×10^5^ cells/ml culture medium with 20 ng/ml GM-CSF in complete RPMI 1640 medium to induce the differentiation of BMDCs. RPMI 1640 medium with GM-CSF was refreshed every 3 days. On day 9, bone marrow derived cells (BMDCs) were harvested by collecting the non-adherent cells after gently swirling the plate and restimulated for 6 h with RPMI 1640 medium alone or cecal microbiota products (CMPs) obtained from mice that received various treatments as indicated in related figure legends. CMPs were prepared by dissolving cecal contents with sterile PBS (100 mg/ml) with vigorous vertex followed by centrifuge at 10,000 rpm for 2 min and filtered through 0.22 μm MillexGP filter unit. The CMP solution was then stored at −20°C and diluted by 40 times using PBS(−) for use. Cells were cultured at 37°C with 5% CO_2_.

### Real-time PCR

RNA purification from sorted MNPs was performed by using the RNeasy Mini Kit (Qiagen). cDNA was obtained by reverse transcription using the High-capacity cDNA Reverse Transcription Kits (ABI). Samples were analyzed by real-time PCR with GoTaq qPCR Master Mix (Promega) on the Applied Biosystem 7900HT Fast machine. Primers were used are *Ifna* forward, 5’-GGACTTTGGATTCCCGCAGGAGAAG-3’; *Ifna* reverse, 5’-GCTGCATCAGACAGCCTTGCAGGTC-3’. *Ifnb* forward, 5’-AACCTCACCTACAGGGCGGACTTCA-3’; *Ifnb* reverse, 5’-TCCCACGTCAATCTTTCCTCTTGCTTT-3’. *Irf7* forward, 5’-CCCCATCTTCGACTTCAGAG-3’; *Irf7* reverse, 5’-AAGGAAGCACTCGATGTCGT-3’. *Isg15* forward, 5’-TGACTGTGAGAGCAAGCAGC-3’; *Isg15* reverse, 5’-CCCCAGCATCTTCACCTTTA-3’. Glyceraldehyde-3-phosphate dehydrogenase (*Gapdh*) forward, 5’-TGAACGGGAAGCTCACTGG-3’; *Gapdh* reverse, 5’-TCCACCACCCTGTTGCTGTA-3’. Expression was normalized to *Gapdh* and presented as relative expression to the control group by the 2^−ΔΔCt^ method.

Total bacterial DNA was extracted from cecal contents using the QIAamp DNA Stool Mini Kit (Qiagen) according to the manufacturer’s protocol. DNA concentration and quality in the extracts was determined by NanoDrop 1000 spectrophotometer (Thermo Scientific). Bacterial groups in cecal samples were measured by real-time PCR using GoTaq qPCR Master Mix (Promega) on Applied Biosystem 7900HT Fast or StepOne Plus™ Real-Time PCR Systems. Primers used for 16S rRNA gene qPCR are as follows ^50–55^: *Firmicutes* forward (Firm934F), 5’-GGAGYATGTGGTTTAATTCGAAGCA-3’; *Firmicutes* reverse (Firm1060R), 5’-AGCTGACGACAACCATGCAC-3’. *Bacteroidetes* forward (Bact934F), 5’-GGARCATGTGGTTTAATTCGATGAT-3’; *Bacteroidetes* reverse (Bact1060R), 5’-AGCTGACGACAACCATGCAG-3’. *Clostridium XIVa* forward, 5’-AAATGACGGTACCTGACTAA-3’; *Clostridium XIVa* reverse, 5’-CTTTGAGTTTCATTCTTGCGAA-3’. *Clostridium sp*. forward, 5’-CACCAAGGCGACGATCAGT-3’; *Clostridium sp*. reverse, 5’-GAGTTTGGGCCGTGTCTCA-3’. *Clostridium coccoides* subgroup forward (CcocF), 5’-AAATGACGGTACCTGACTAA-3’; *Clostridium coccoides* subgroup reverse (CcocR), 5’-CTTTGAGTTTCATTCTTGCGAA-3’. *Clostridium coccoides-Eubacteria rectale* group forward (ClEubF), 5’-CGGTACCTGACTAAGAAGC-3’; *Clostridium coccoides-Eubacteria rectale* group reverse (ClEubR), 5’-AGTTTYATTCTTGCGAACG-3’. *ASF500* forward (500-183F), 5’-GTCGCATGGCACTGGACATC-3’; *ASF500* reverse (500-445R), 5’-CCTCAGGTACCGTCACTTGCTTC-3’. *ASF360* forward (360-81F), 5’-CTTCGGTGATGACGCTGG-3’; *ASF360* reverse (360-189R), 5’-GCAATAGCCATGCAGCTATTGTTG-3’. All bacteria forward (UnivF), 5’-TCCTACGGGAGGCAGCAGT-3’; All bacteria reverse (UnivR), 5’-GACTACCAGGGTATCTAATCCTGTT-3’. Expression was normalized to all bacterial DNA, and relative expression levels were calculated relatively to the vehicle control group by the 2^−ΔΔCt^ method.

### 16S rRNA gene sequencing

Aliquots for sequencing of the 16S rRNA gene were first amplified with the V3-V4 region primers 341F (5’-CCTACGGGAGGCAGCAG-3’) and 518R (5’-ATTACCGCGGCTGCTGG-3’) as described previously ^56^. A reagent-only control (DNA extraction kit blank) and a mock bacterial community (HM-782D, BEI Resources, ATCC, Manassas, VA) were also prepared in the same manner. A single library pool was compiled using equimolar concentrations of DNA as measured using a fluorometric assay (Qubit dsDNA Broad-Range Assay kit, Invitrogen, UK). The Illumina MiSeq platform (Illumina, CA) was used for sequencing (Edinburgh Genomics, United Kingdom), using V2 chemistry and producing 250 bp paired-end reads. Using the mock bacterial community data, the sequencing error rate was calculated as 0.01%. The raw sequence reads, with primers removed, are publicly available via the NCBI Sequence Read Archive (SRA) under accession number PRJNA564944.

### 16sRNA sequencing analysis

For the raw 16s RNA data in FASTQ format, amplicon primers were first removed to prevent false positive detection of chimeras using the cutadapt plugin ^57^. 10,000 sequences were sampled at random and the qualities at each base position were examined for determining the parameters of the denoising process. The paired-end reads were further trimmed, filtered, denoised and merged using the DADA2 plugin ^58^ through the Wales supercomputer portal. A naïve Bayes classifier within QIIME2 v2019.10 ^59^ was trained against the Silva v132 database (https://www.arb-silva.de/) on the amplified region. And a machine learning Python library scikit-learn ^60^ was used to classify operational taxonomic units (OTUs) based on 100% sequence identity. A total of 7 taxonomic levels were used for 16S rRNA datasets. For measuring diversity, we generated de novo phylogenetic trees through multiple sequence alignment, masking, tree building and rooting using the MAFFT program ^61^. Percent abundance of taxa was determined by calculating it as a proportion of the total read count across all samples. For instance, the percent abundance of the *i*-th taxa in the *j-th* sample is computed as

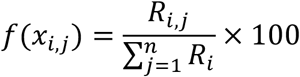

where *n* is the total number of samples and *R* represents read counts of each taxa.

The OTU table was rarefied across samples to the 90% of the lowest sample depth with a random seed set as 123 to eliminate the bias caused by the different sample sizes. For the overall bacterial community within samples, alpha diversity estimators including the observed species, the Chao1 index, the Shannon diversity, and the Inverse Simpson (Invsimpson) were calculated using Phyloseq ^62^. The alpha diversity estimates were compared between three groups using non-parametric Kruskal-Wallis test along with Dunn’s multiple comparison correction. Beta-diversity between samples was computed using the unweighted UniFrac distances ^63^.

### Statistical analysis

All data were expressed as mean ± SD except that in Fig 8A, where the data were expressed as mean ± SEM as indicated in the figure legends. Statistical significance between two groups was examined by unpaired Student’s *t*-test, while the analysis of variance (ANOVA) with post-hoc Holm-Sidak’s multiple comparisons test was used to evaluate multiple groups. Statistical work was performed using Prism 8 software (GraphPad) and a *P* value of less than 0.05 was considered as significance.

## Acknowledgments

We thank P. Ghazal for helpful discussions and support; F Rossi, S Johnston, W Ramsay and M Pattison at the University of Edinburgh QMRI and SCRM flow cytometry facilities for cell sorting and analysis. We acknowledge the support of the Supercomputing Wales project, which is part-funded by the European Regional Development Fund (ERDF) via Welsh Government. This work was supported in part by Medical Research Council (MRC) UK (MR/R008167/1 to CY; MR/K013386/1 to AGR; MR/P008887/1 to DJM; MR/M011445/1 to VOD), Cancer Research UK (C63480/A25246 to CY), the Wellcome Trust Investigator Award (106122 to RMM), the Rainin Trust Award (13H6 to RMM), European Research Council (LipidArrays to VOD), the Pathological Society of Great Britain and Ireland (MJA), Core Research for Evolutional Science and Technology Program from the Japan Agency for Medical Research and Development (SN) and the collaborative grant to the Kyoto University from Ono Pharmaceuticals (SN). SC received a PhD studentship by The University of Edinburgh.

## Author contributions

SC and MG designed and performed experiments and analyzed the data with assistance by CTB, DJS, AA, RAO, XZ, AM, HXY and CY. JP, MG, YZ and RA performed microbiome 16S rRNA sequencing and data analysis. GTH and MJA analyzed intestine histology. VT, LD and VOD performed lipidomic analysis. XFL, BZQ, JKJS and RMB contributed to critical reagents and the generation of transgenic animal lines. SV, DJM, MJA, JPI, SMA, SN, RMM, AGR and SEH provided technical expertise, essential reagents and transgenic mice, advised on experiment design and data analysis, and edited and critiqued the manuscript. CY conceived this project, supervised the research and wrote the manuscript.

