## Supplementary material for "Prostaglandin E_2_ promotes intestinal inflammation via inhibiting microbiota-dependent regulatory T cells": Supplemetal materials

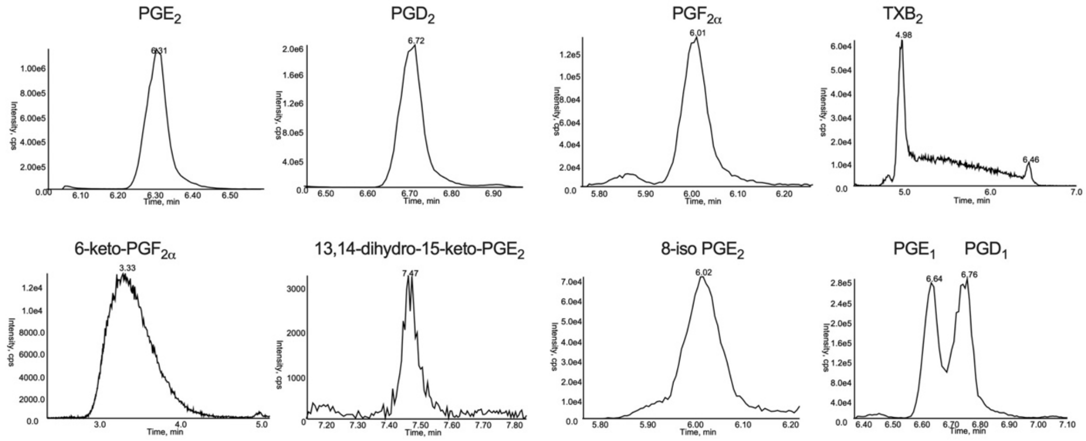


**Supplementary Figure 1. Detection of PGs in intestinal tissues.** Representative chromatogram for each detectable prostaglandins (PGs) and their metabolites in small intestines and colons from mice administered with indomethacin or vehicle control in drinking water for 5 days. Levels of $PGD_3$, $PGE_3$, 13,14-dihydro-15-keto $PGF_{2\alpha}$, 8-iso-15-keto $PGF_{2\alpha}$ and 13,14-dihydro-15-keto $PGD_2$ in both small intestine and colon tissues were undetected.

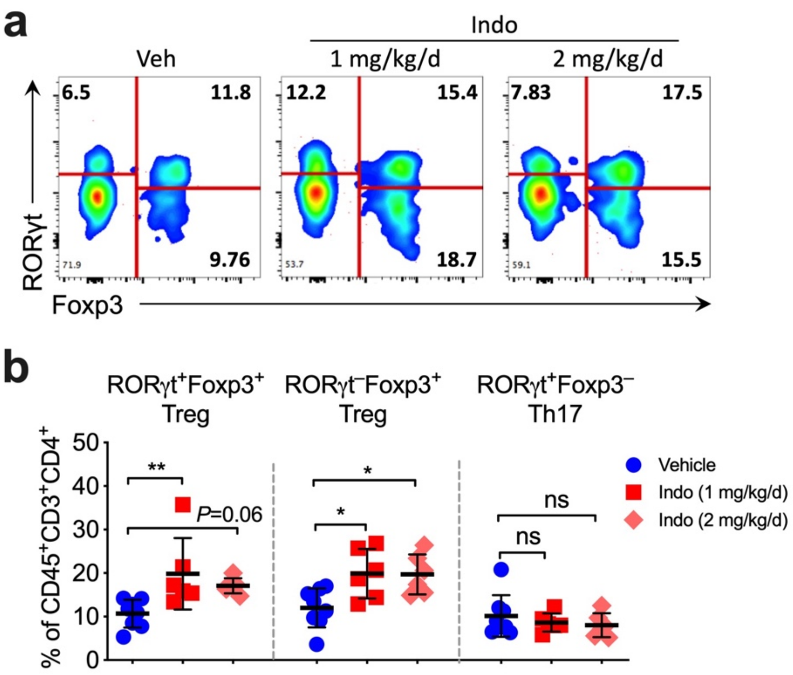


**Supplementary Figure 2. Effects of low doses of indomethacin on colonic Tregs.** Representative flow cytometry plots (**a**) and percentages (**b**) of RORγt$^+$Foxp3$^+$ Tregs, RORγt$^-$Foxp3$^+$ Tregs, and RORγt$^+$Foxp3$^-$ Th17 cells gated on live CD4$^+$ T cells in colon lamina propria of mice treated with vehicle (n=8) or different doses of indomethacin (Indo, 1 or 2 mg/kg/d, each n=6) for 2 weeks. Each scatter dot plot in bar graphs represents data from one mouse. Data shown as mean ± SD are pooled from two independent experiments and analysed by ANOVA with post-hoc Holm-Sidak's multiple comparisons test. **P*<0. 05, ***P*<0.01, ns=not significant.

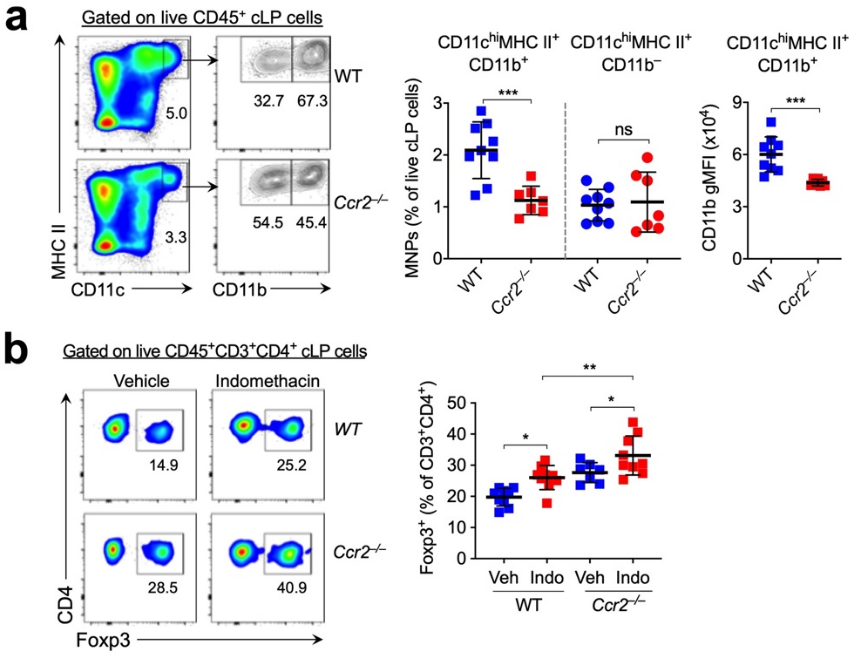


**Supplementary Figure 3. Effects of CCR2 deficiency on indomethacin-dependent regulation of colonic Tregs. (a)** Representative flow cytometry dot-plots (left) and percentages of colon lamina propria (cLP) CD11c$^+$MHC II$^+$CD11b$^+$ and CD11c$^+$MHC II$^+$CD11b$^-$ MNPs (middle) in WT and *Ccr2*$^{-/-}$ mice (n=9 each group). CD11b gMFI of CD11c$^+$MHC II$^+$CD11b$^+$ MNPs are also shown (right). **(b)** Representative flow cytometry dot-plots (left) and percentages of cLP Foxp3$^+$ Tregs among CD3$^+$CD4$^+$ T cells in WT and *Ccr2*$^{-/-}$ mice treated with indomethacin (Indo) or vehicle control (Veh) (n=7-9). Each scatter dot plot in bar graphs represents data from one mouse. Data shown as mean ± SD are pooled from two independent experiments and analysed by two-tailed unpaired student *t*-test (**a**) or ANOVA with post-hoc Holm-Sidak's multiple comparisons test (**b**). *$P$<0.05, **$P$<0.01, ***$P$<0.001, and ns=not significant.
